# PEN receptor GPR83 in anxiety-like behaviors: differential regulation in global vs amygdalar knockdown

**DOI:** 10.1101/2021.05.13.443933

**Authors:** Amanda K. Fakira, Lindsay M. Lueptow, Nikita A. Trimbake, Lakshmi A. Devi

## Abstract

Anxiety disorders are prevalent across the United States and result in a large personal and societal burden. Currently, numerous therapeutic and pharmaceutical treatment options exist. However, drugs to classical receptor targets have shown limited efficacy and often come with unpleasant side effects, highlighting the need to identify novel targets involved in the etiology and treatment of anxiety disorders. GPR83, a recently deorphanized receptor activated by the abundant neuropeptide PEN, has also been identified as a glucocorticoid regulated receptor (and named GIR) suggesting that this receptor may be involved in stress-responses that underlie anxiety. Consistent with this, GPR83 null mice have been found to be resistant to stress-induced anxiety. However, studies examining the role of GPR83 within specific brain regions or potential sex differences have been lacking. In this study, we investigate anxiety-related behaviors in male and female mice with global knockout and following local GPR83 knockdown in female mice. We find that a global knockdown of GPR83 has minimal impact on anxiety-like behaviors in female mice and a decrease in anxiety-related behaviors in male mice. In contrast, a local GPR83 knockdown in the basolateral amygdala leads to more anxiety-related behaviors in female mice. Local GPR83 knockdown in the central amygdala or nucleus accumbens showed no significant effect on anxiety-related behaviors. Finally, dexamethasone administration leads to a significant decrease in receptor expression in the amygdala and nucleus accumbens of female mice. Together, our studies uncover a significant, but divergent role for GPR83 in different brain regions in the regulation of anxiety-related behaviors, which is furthermore dependent on sex.

## Introduction

Anxiety disorders manifest in a variety of symptoms, but often involve excessive and/or persistent worry and fear that is considered maladaptive and intensifies over time. These disorders occur in approximately 20% of the population, and therefore represent a significant societal and economic burden. Anxiety disorders also occur at a higher rate in females compared to males (Palanza, 2001; Boivin et al., 2017; Zuloaga et al., 2020) (see also www.nih.nimh.gov), and females respond differentially to anxiolytic drugs (Palanza, 2001) suggesting differences in the etiologies and/or underyling circuitry for anxiety between males and females. There are several neurotransmitter systems such as the γ-aminobutyric acid (GABA)-ergic and serotinergic systems, as well as neuropeptide systems including neuropeptide Y, cholecystokinin, corticotrophin releasing factor, and substance P that are targets for anxiety disorders (Griebel and Holmes, 2013; Murrough et al., 2015). Despite decades of continued research into the treatment of anxiety by targeting these major neurotransmitter systems, little progress has been made in the development of more efficacious treatments. This has led to a shift in research focus to other lesser known neurotransmitter and peptide systems with the intent of identifying the next generation of anxiolytics. One underutilized source of novel therapuetics is the pool of orphan G protein-coupled receptors (GPCRs) whose endogenous ligands are beginning to be explored.

Studies focussing on deorphanizing hypothalamic orphan GPCRs led to the identification of the abundant neuropeptide PEN as an endogenous ligand for the orphan receptor GPR83 (Gomes et al., 2016; Foster et al., 2019; Parobchak et al., 2020). The PEN peptide is a product of the cleavage of the proSAAS precursor (Fricker et al., 2000; Mzhavia et al., 2002), and has been implicated in a number of neurologic functions and disorders including feeding, reward, and Alzheimer’s disease (Wei et al., 2004; Wardman et al., 2011; Hoshino et al., 2014; Wang et al., 2016a; Berezniuk et al., 2017). In fact, proSAAS and its peptide products have also been implicated in anxiety-related behavior (Wei et al., 2004; Morgan et al., 2010; Bobeck et al., 2017). Specifically for GPR83, it was found that the glucocorticoid receptor agonist, dexamethasone, regulates its expression in immune cells and the brain (Harrigan et al., 1989, 1991; Adams et al., 2003) suggesting that activation of stress-responses regulates GPR83 expression. Finally, a study examining behaviors of mice lacking GPR83 noted that they were resilient to stress-induced anxiety (Vollmer et al., 2013). These data have suggested a role for GPR83 in modulating anxiety-related behaviors (Lueptow et al., 2018; Mack et al., 2019).

In this study, we sought to directly investigate the role of GPR83 in anxiety-related behaviors, with a specific focus on female mice who have been previously overlooked, and furthermore, to determine the extent to which GPR83 expression in the amygdala subnuclei and nucleus accumbens contribute to these behaviors. To date, no small molecule agonists or antagonists for this receptor have been identified, therefore we used a combination of GPR83 global knockout (KO) animals and GPR83 shRNA mediated local knockdown (KD) in the basolateral amygdala (BLA), central nucleus of the amygdala (CeA) and nucleus accumbens (NAc) to study the role of this receptor in anxiety-related behaviors. These brain regions play a role in anxiety, display significant GPR83 expression, and form circuits that encode positive and negative affective valence (Stuber et al., 2011; Tye et al., 2011; Janak and Tye, 2015a; Namburi et al., 2015; Tovote et al., 2015; Beyeler et al., 2016; Lueptow et al., 2018; Fakira et al., 2019). Our initial behavioral studies also included both male and female subjects to determine whether GPR83 plays a differential role in anxiety-related behaviors between the two sexes. We found that the knockdown of GPR83 in the basolateral amygdala of female mice resulted in increased anxiety-related behaviors and that dexamethasone administration led to sex-specific regulation of GPR83 expression supporting a role for this receptor in modulating anxiety-related behaviors which are dependent on sex.

## Materials and Methods

### Animals

GPR83/eGFP (Rockefeller University, NY, NY), GPR83 KO and C57BL/6J (Jackson Labs, Bar Harbor ME) male and female mice (8-12 weeks) were maintained on a 12hr light/dark cycle with water and food ad libitum. GPR83/eGFP BAC transgenic mice were generated by the GENSAT project at Rockefeller University. The coding sequence for enhanced green fluorescent protein (eGFP) followed by a polyadenylation signal was inserted into a bacterial artificial chromosome (BAC) at the ATG transcription codon of GPR83. Therefore, cells that express GPR83 mRNA also express eGFP. GPR83 knockout (Jackson Labs, Bar Harbor, ME) mice, generated previously (Lu et al., 2007), lack GPR83 protein and mRNA (Gomes et al., 2016; Fakira et al., 2019). Animal protocols were approved by the IACUC at Icahn School of Medicine at Mount Sinai, according to NIH’s Guide for the Care and Use of Laboratory Animals. The number of animals of each sex per group for each experiment is indicated on the individual figure legends.

### Elevated Plus Maze and Open Field Assay

One week prior to testing mice were handled for five minutes a day for three to four days. On the day of testing, mice were habituated to the testing room 1-hour before open field testing followed by the elevated plus maze four hours later. The open field consists of a 40 × 40 × 40 cm box made of white plastic material. Mice explored the open field for 30 minutes under red light illumination. The amount of time spent in the center of the open field was tracked for the first 5 minutes and the distance traveled was tracked for 20 minutes. The elevated plus maze consisted of two open and two closed arms (12 × 50 cm each) on a pedestal 60 cm above the floor. Mice explored the maze for 5 min under red light. The amount of time spent in each arm was tracked and analyzed using Noldus EthoVision XT. Data are presented as center time (s), distance traveled (cm), and open arm time (%; open arm time/ (open arm + closed arm time)). Vaginal swabs from the female mice were collected and visualized immediately following behavioral testing as described previously (McLean et al., 2012). Images were collected from the vaginal smears from each animal. The images were later visualized by two blinded investigators who categorized them as proestrus, estrous, metestrus and diestrus by the presence of nucleated epithelial cells, cornified epithelial cells and leukocytes. During metestrus and diestrus leukocytes predominate while, proestrus and estrus is characterized by nucleated and cornified epithelial cells. For analysis, female mice were grouped together as mice in estrus/proestrus, when follicle stimulating hormone, estradiol, luteinizing hormone, and prolactin levels are high. This group of mice are referred to as oestrus throughout the paper. The mice in diestrus and metestrus, when progesterone levels predominate, were also grouped together and are referred to as diestrus throughout the paper (Miller and Takahashi, 2014).

### Immunofluorescence and Confocal Imaging

GPR83/eGFP mice were perfused with 4% paraformaldehyde in phosphate buffered saline pH 7.4. The brains were removed and post fixed in 4% paraformaldehyde in phosphate buffered saline overnight. Brains were rinsed 3 times in phosphate buffered saline and 50 μM coronal brain slices were obtained using a vibratome (Leica VT1000, Buffalo Grove, Il), without embedding the tissue. To visually enhance eGFP expression, immunohistochemical analysis was carried out using chicken anti-GFP (1:1000) as the primary antibody (Aves Labs, Tigard, OR) and anti-chicken 488 (1:1000) as the secondary antibody (Molecular Probes, Eugene, OR). In addition, brain slices were co-stained for parvalbumin (1:250; ThermoFischer Scientific, Rockford, Il) overnight followed by anti-sheep 568 (1:500) secondary antibody. Confocal microscopy was performed in the Microscopy CoRE at the Icahn School of Medicine at Mount Sinai. Confocal z-stack images were taken on a Zeiss LSM 780 microscope and processed using Zeiss software. Confocal images of the amygdala were taken from regions from coronal sections between Bregma −0.58 and −2.06 mm (Paxinos and Franklin, 2012).

### Dexamethasone Treatment and Quantitative PCR

Male and female mice were injected with dexamethasone (5mg/kg; i.p.). Three hours later amygdala and NAc punches were collected for qPCR analysis. Total cellular RNA was extracted from amygdala and NAc punches using Qiazol reagent and the RNAeasy Midi kit (QIAGEN, Valencia, CA). Total RNA was reverse transcribed into cDNA using VILO master mix (Invitrogen, Carlsbad, CA). qPCR was performed in triplicate aliquots from each individual animal with Power SYBR Green PCR master mix (ThermoFisher, Waltham, MA), 25 ng of cDNA and 0.5 μM of primers using an ABI Prism 7900HT (Thermo Fisher, Waltham, MA) in the qPCR CoRE at Icahn School of Medicine at Mount Sinai. Primer sequences for GPR83, proSAAS and GAPDH are the same as used previously (Fakira et al., 2019). The primer sequences used for qPCR are: GAPDH Forward: 5’-TGAAGGTCGGTGTGAACG Reverse: 5’-CAATCTCCACTTTGCCACTG, GPR83 Forward: 5’-GCAGTGAGATGCTTGGGTTC Reverse: 5’-CCCACCAATAGTATGGCTCA and proSAAS: Forward: 5’-AGTGTATGATGATGGCCC Reverse: 5’-CCCTAGCAAGTACCTCAG. The CT values for the technical replicates were averaged and analysis performed using the CT method and normalized to saline controls. In some cases, qPCR reactions were repeated to determine the reliability of the primers and RNA samples.

### GPR83 shRNA and surgeries

Three weeks prior to behavioral testing, a craniotomy was performed under isoflurane anesthesia and 0.5 μL of lentiviral GPR83 shRNA or control shRNA particles (10^9^; Sigma Mission Lentiviral Transduction Particles, St. Louis, MO) were infused into the NAc (A/P: +1.5, Lat: +/−1.6, D/V:−4.4), BLA (A/P: −1.1, Lat: +/− 3.2, D/V:−5.1) or CeA (A/P: −1.0, Lat: +/− 2.8, D/V:−4.9). The GPR83 shRNA targeted the sequence 5’-CCATGAGCAGTACTTGTTATA-3’, an exonic region of the gene. A nucleotide BLAST of this sequence produces three alignments with E values of 0.003 that correspond to GPR83 variants; other alignments have E values greater than 40, indicating that this sequence has few off targets.

### Data Analysis

Data are presented as mean ± SEM. Data were analyzed using Student’s t-test or Two-way ANOVA with Bonferroni’s Multiple Comparison post-hoc tests using GraphPad Prism 8.0 software (San Diego, CA). The number of animals/group for each experiment is indicated on the individual figure legends.

## Results

### Analysis of anxiety-related behaviors in male and female GPR83 KO mice

Anxiety-related behaviors in GPR83 WT and KO mice were analyzed using the elevated plus maze (Figure 1A) and open field tests (Figure 1D). GPR83 KO mice spent more time in the open arms of the elevated plus maze (EPM) compared to wild type mice, indicating lower anxiety-like behavior in KO mice; however, there was no difference in the frequency to enter the open arms (Figure 1B and C). In the open field assay, there was no difference in the amount of time GPR83 KO mice spent in the center of the open field; however, there was a significant increase in the frequency that they entered the center region, also an indication of lower anxiety-like behavior (Figure 1E and F). Overall, these differences are unlikely to be due to changes in overall motor activity of GPR83 KO mice, since there were no differences in locomotor activity in the open field (Figure 1G). Together, these results show that global loss of GPR83 produces a decrease in anxiety levels.

**Figure 1:**
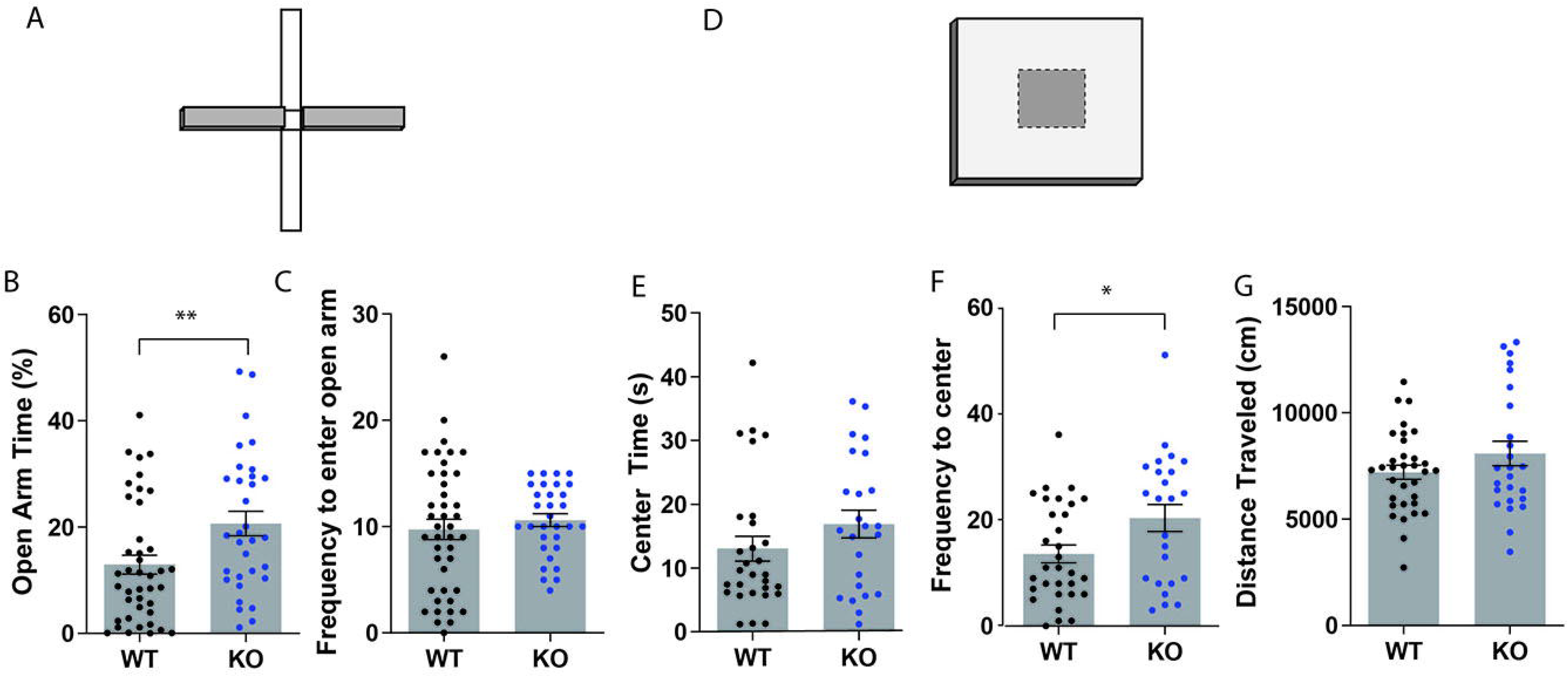
Mice lacking GPR83 have a decrease in anxiety-related behaviors. Wild type (WT) and GPR83 knockout (KO) mice were screened on the elevated plus maze **(A)** and the amount of time spent on the open arms **(B)** and frequency to enter the open arms **(C)** measured. WT and GPR83 KO mice were screened in an open field assay **(D)** and the amount of time spent in the center **(E)**, frequency to enter to center **(F)** and distance traveled (**G**) measured. Data are represented as mean ± SEM and analyzed using Student’s T-test, % = open arm time/ (open arm + closed arm time), *p<0.05; **p<0.01; WT, n= 41; GPR83 KO, n=32.

The data were further analyzed to determine the extent to which sex differences might contribute to anxiety-related behaviors in the global GPR83 KO mice In the EPM test, when separated by sex, GPR83 KO mice exhibited a significant effect on open arm time, with no interactions between sex and GPR83 KO genotype (Figure 2B; 2-way ANOVA; Interaction F_(1,70)_=0.11, p=0.7436; GPR83 KO F_(1,70)_=5.02, p=0.0282; Sex F_(1,70)_=1.14, p=0.2891). In addition, Bonferroni’s post-hoc analysis did not indicate any differences between groups; however, male GPR83 KO mice spent more time in the open arm of the elevated plus maze compared to wild type when analyzed by Student’s T-test (Figure 2B; *p<0.05). Neither analysis revealed any differences in the frequency to enter the open arm for either sex (Figure 2C, 2-way ANOVA; Interaction F_(1,70)_=0.57, p=0.4525; GPR83 genotype F_(1,70)_=1.74, p=0.1918; Sex F_(1,70)_=0.98, p=0.3258). Direct comparison of open arm time in wild type males and females indicates that there is a tendency for female mice to spend more time in the open arm compared to males (Figure 2B; Student’s t-test @ p=0.0872), indicating that female mice may be less anxious compared to males overall.

**Figure 2:**
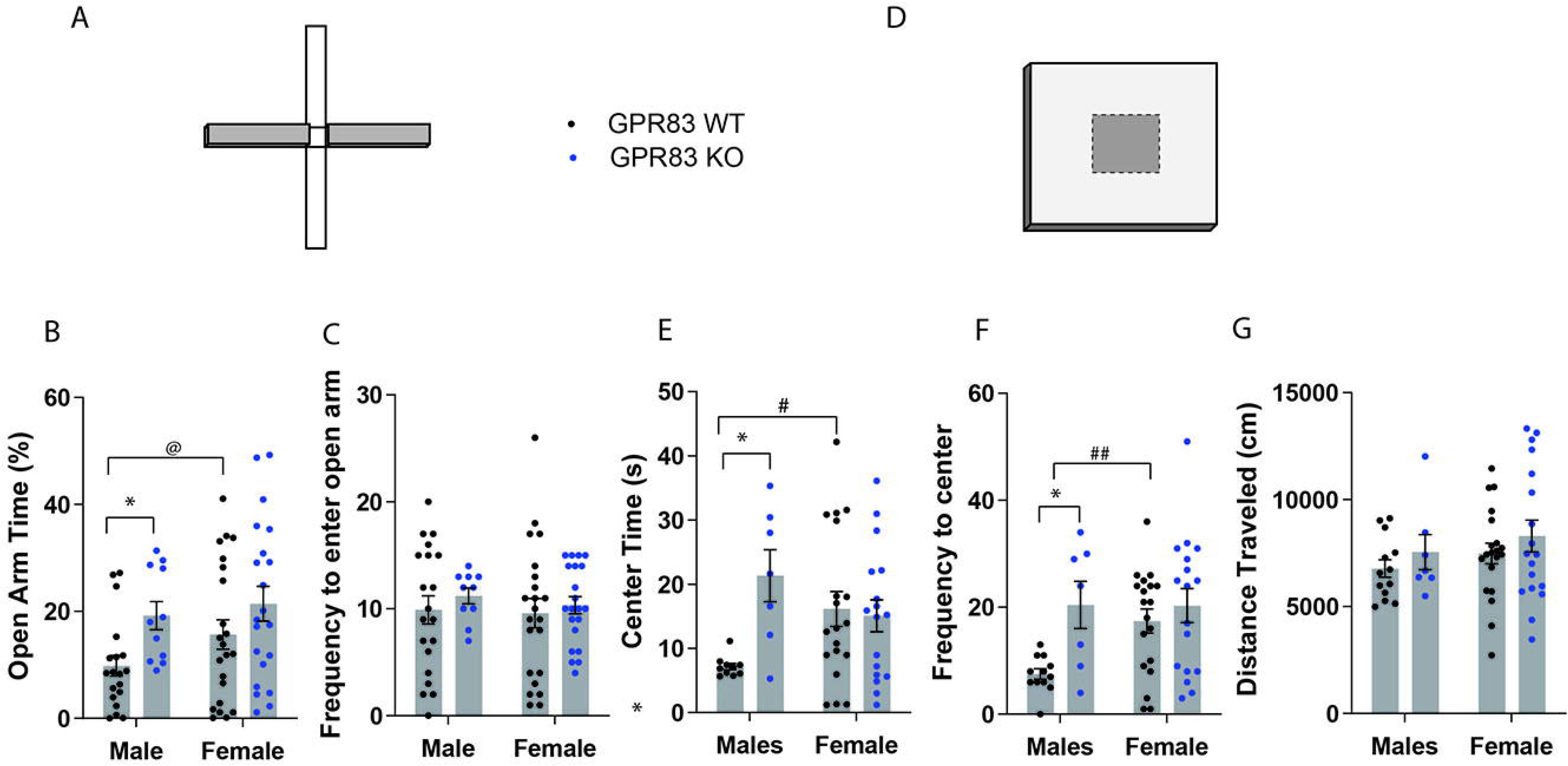
Sex-differences in GPR83-mediated regulation of anxiety-related behaviors. Sex-dependent analysis of WT and GPR83 KO mice on the elevated plus maze **(A)** measuring open arm time **(B)** and frequency to enter the open arm **(C)**. Sex-dependent analysis of WT and GPR83 KO mice in the open field assay **(D)** measuring center time **(E)**, frequency to enter the center **(F)** and distance traveled **(G)**. Data are represented as mean ± SEM and analyzed using 2-way ANOVA following Bonferroni’s post-hoc test (*p<0.05) and Student’s T-test, %= open arm time/ (open arm + closed arm time), @ p=0.0872; #p<0.05, ##p<0.01; WT males, n=19, GPR83 KO males, n= 11; WT females, n=22, GPR83 KO females, n=21.

In the open field test the sex-dependent analysis of anxiety-related behavior revealed that male GPR83 KO mice spent significantly more time in the center compared to wild type males, an effect that was not seen when comparing female GPR83 KO mice with wild type females (Figure 2E; 2-way ANOVA; Interaction F_(1,51)_=6.34, p=0.0150; GPR83 KO F_(1,51)_=4.67, p=0.0355; Sex F_(1,51)_=0.15, p=0.7044; Bonferroni’s post-hoc test Males: WT vs GPR83 KO, *p<0.05). Moreover, analysis of frequency to enter the center indicates a similar effect in that male GPR83 KO mice entered the center of the open field more frequently than male wild type mice while no differences were seen between female GPR83 KO mice and female wild type mice (Figure 2F; 2-way ANOVA; Interaction F_(1,51)_=2.84, p=0.0980; GPR83 F_(1,51)_=7.01, p=0.0.0107; Sex F_(1,51)_=2.69, p=0.1071; Bonferroni’s post-hoc test Males: WT vs GPR83 KO, *p<0.05). Analysis of baseline anxiety differences between wild type male and female mice indicate that female mice display significantly less anxiety-related behaviors levels, spending more time in the center (#p<0.05) and entering the center of the open field more frequently (## p<0.01) than males (Figure 2E and F). In addition, we find no effect of GPR83 KO on locomotor activity levels even when segregated by sex (Figure 2G; 2-way ANOVA; Interaction F_(1,54)_=0.001201, p=0.9725; GPR83 F_(1,54)_=1.137, p=0.0.2911; Sex F_(1,54)_=1.341, p=0.2520). Overall, these data indicate that lack of GPR83 produces a decrease in anxiety-related behaviors that is more pronounced in male mice, likely due to their higher levels of baseline anxiety compared to female mice.

### Cell-type specific expression of GPR83 expression in the amygdala

The amygdala is a brain region well known to play a role in anxiety-related behaviors (Gilpin et al., 2015; Janak and Tye, 2015b; Tovote et al., 2015). Within the basolateral amygdala (BLA), parvalbumin cells (PV^+^) are the largest population of GABAergic inhibitory interneurons (McDonald and Mascagni, 2001), directly influencing output of primary excitatory neurons, and these cells have been directly implicated in anxiety-like behavior (Urakawa et al., 2013; Babaev et al., 2018). In addition, PV^+^ neurons within the central amygdala (CeA), have been implicated in anxiety trait (Ravenelle et al., 2014), as well as opioid withdrawal-induced negative affect, including anxiety-like behavior (Wang et al., 2016b). *In situ* hybridization data from the Allen Mouse Brain Atlas indicates that GPR83 is expressed in both the BLA and CeA (Figure 3A and B). Therefore, we sought to examine the co-expression of PV^+^ and GPR83^+^ cells in this brain region using GPR83/GFP BAC transgenic mice. These mice express eGFP under control of the GPR83 promotor; therefore all cells that express the receptor will also express eGFP. We find that GPR83 is expressed throughout the amygdala (Figure 3C) with higher expression in the BLA and the CeA (Fig 3C and D). Higher magnification images demonstrate that the eGFP positive cells have a neuronal morphology (Fig 3E). Subsequent co-staining with parvalbumin indicates that some of the GPR83 positive cells express parvalbumin and, therefore are GABAergic neurons (Figure 3F-H).

**Figure 3:**
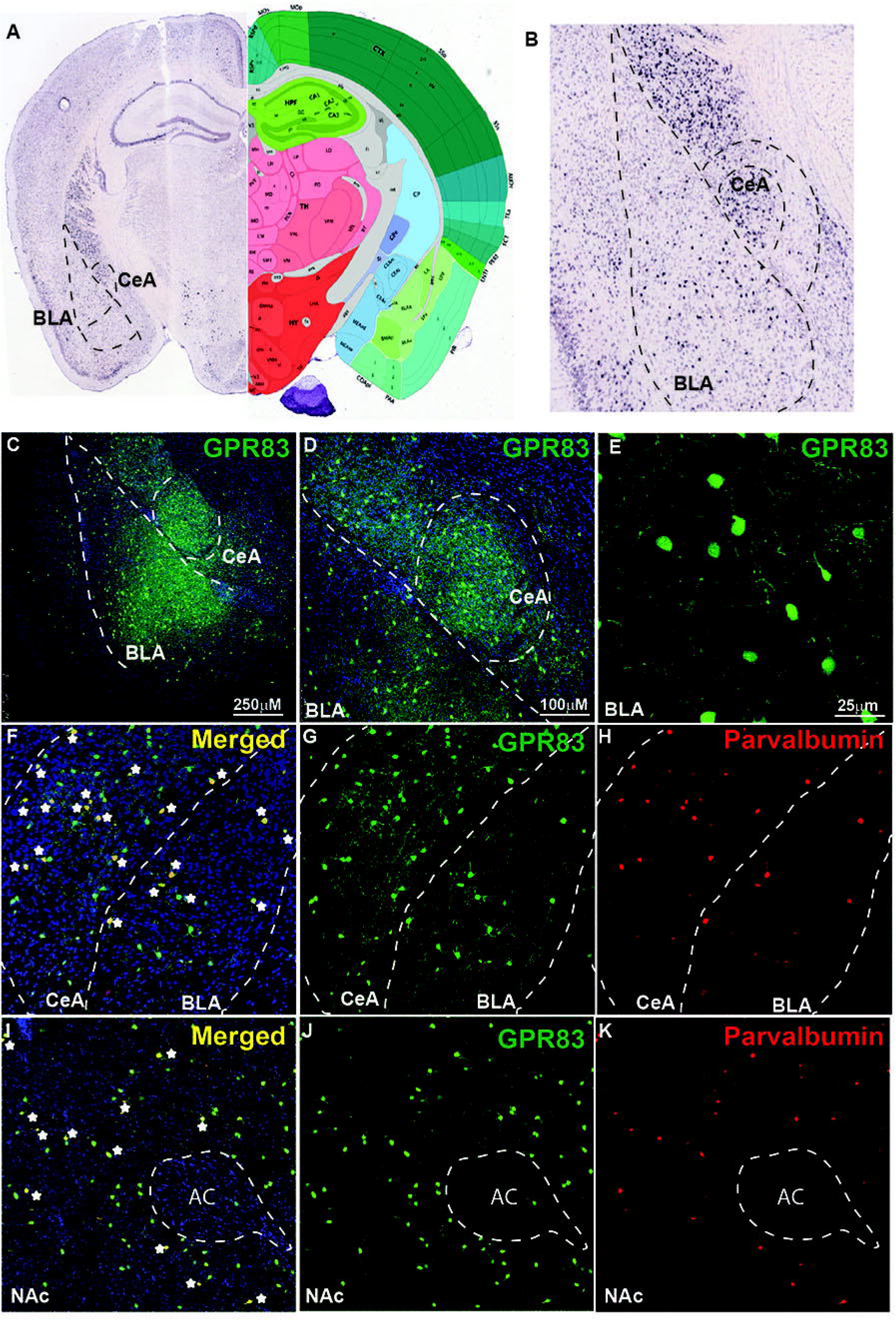
GPR83 expression in the basolateral and central nucleus of the amygdala. **(A)** In situ hybridization image for GPR83 from the Allen Mouse Brain Atlas (Allen Institute. © 2015 Allen Institute for Brain Science. Allen Brain Atlas API) (left) and corresponding brain atlas image (right). **(B)** Enlarged image of BLA and CeA shown in **(A)**. Low magnification image of GPR83 (green) expression in the BLA and CeA **(C-D)** using GPR83-GFP reporter mice from Gensat. **(E)** Higher magnification image showing GPR83 expression in neurons in the amygdala. **(F)** Co-localization (yellow, stars) of GPR83 (green, **G**) and parvalbumin (red, **H**) in the BLA and CeA. **(I)** Co-localization (yellow, stars) of GPR83 (green, **J**) and parvalbumin (red, **K**) in the NAc. AC, anterior commissure.

The nucleus accumbens (NAc) is another brain region that contains a high concentration of GPR83 positive cells, and has been strongly implicated in vulnerability and resilience responses to stress (Zhu et al., 2017) as well as anxiety-like behaviors (Xiao et al., 2020). In contrast to the amygdala, we have recently reported that GPR83 is primarily expressed in cholinergic interneurons in the NAc (Fakira et al., 2019). However, a small percentage of neurons were not characterized but recent studies demonstrated that PV^+^ neurons in the striatum indeed express GPR83 (Enterría-Morales et al., 2020). Because PV^+^ neurons in the NAc have been specifically implicated in anxiety-like approach behaviors, we also characterized PV^+^ and GPR83^+^ co-expression in the NAc. We find that a small population of GPR83 positive cells in the NAc also express parvalbumin suggesting the presence of this receptor in some GABAergic neurons (Fig 3I-K).

### Divergent regulation of GPR83 in male and female mice following acute dexamethasone administration

Studies have shown that GPR83 expression is regulated by the glucocorticoid agonist dexamethasone (Harrigan et al., 1989; Adams et al., 2003) suggesting a role for GPR83 in the stress response. Because of this known association with the glucocorticoid system, we used a single dose of dexamethasone to assess its effects on GPR83 expression, in both amygdala and the nucleus accumbens (NAc) of male and female mice. As a control, we examined proSAAS expression, since proSAAS is the precursor to the endogenous ligand for GPR83, PEN. We find that although dexamethasone administration has no effect on GPR83 expression in the amygdala of male mice and a decrease in expression in female mice (Figure 4A and B). In contrast, in the NAc, dexamethasone administration leads to an increase in GPR83 expression in male mice and a decrease in expression in female mice (Figure 4C and D). ProSAAS expression was unchanged by dexamethasone administration in either sex or two brain regions tested (Fig 4A-D). These results suggest that GPR83 expression is regulated by glucocorticoids in a region-specific and sex-dependent manner.

**Figure 4:**
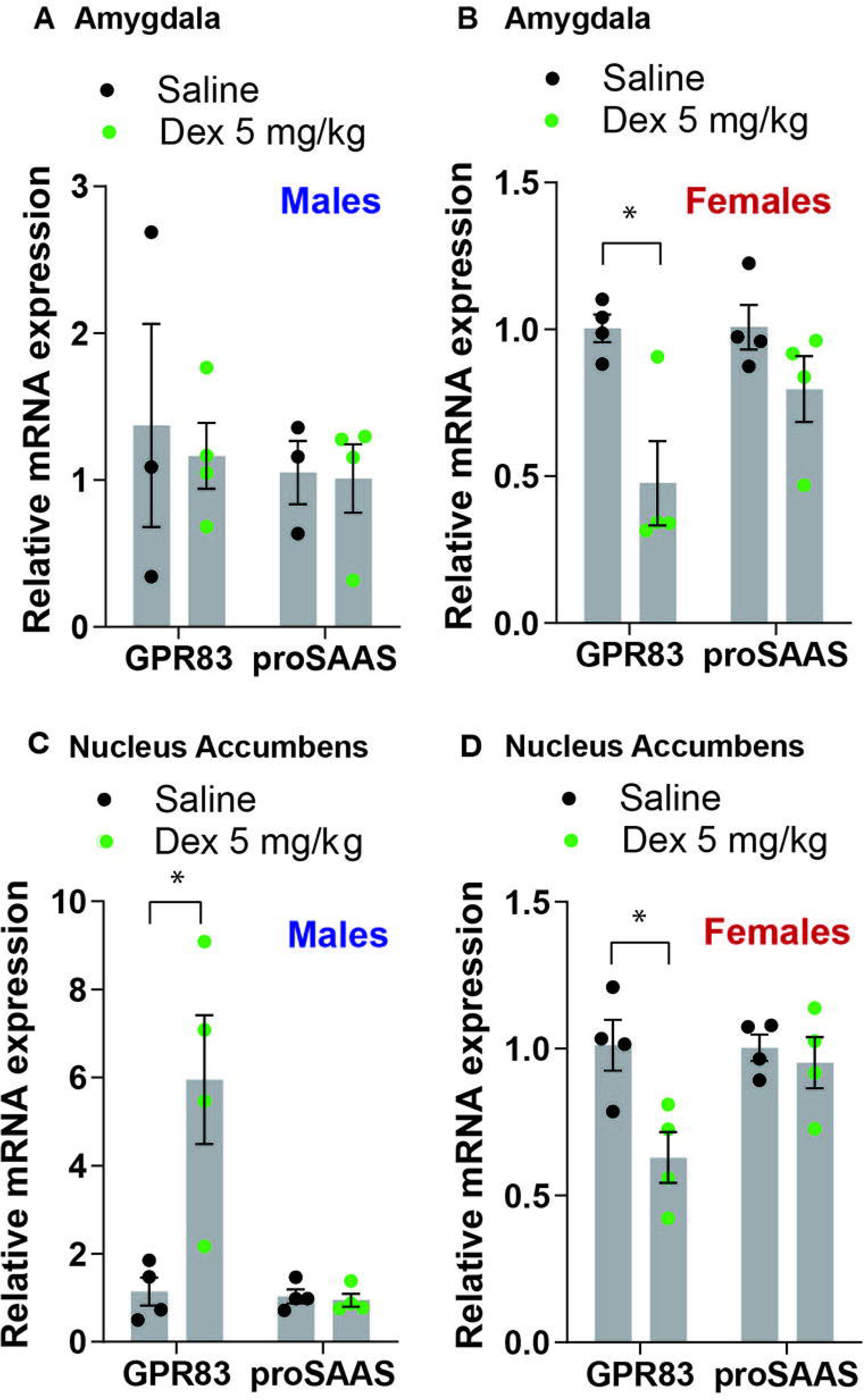
GPR83 expression is regulated by dexamethasone in the amygdala and nucleus accumbens in a sex-dependent manner. **(A and B)** Administration of dexamethasone (5 mg/kg) decreases expression of GPR83 in the amygdala of female mice but not males with no effects on proSAAS expression. **(C and D)** In the NAc, administration of dexamethasone increases expression of GPR83 in males and decreases the expression in females. The proSAAS expression is unchanged in all cases. Data are represented as mean ± SEM and analyzed using Student’s T-test, *p<0.05, n=3-4 per group.

### The effect of GPR83 knockdown in the BLA, CeA and NAc on anxiety-related behaviors

Since glucocorticoids, which are typically released during stressful events that produce anxiety, induce a decrease in GPR83 expression in the amygdala and NAc of female mice we sought to determine the effect of local GPR83 knockdown (GPR83 KD) in the BLA, CeA, and NAc on anxiety-related behaviors in female mice. The knockdown of GPR83 expression was accomplished by administration of GPR83 shRNA lentiviral particles (0.5 μl @10^9^ particles/μl) into the BLA, CeA or NAc of female mice (Figure 5A, G and M) and anxiety-related behaviors analyzed using the elevated plus maze and open field tests and compared to mice that were administered with control virus. In a previous study we showed that this paradigm of lentiviral GPR83 shRNA administration produces a ~50% knockdown compared to control virus (Fakira et al., 2019). We find that local GPR83 KD in the BLA resulted in a decrease in the amount of time spent (***p<0.001) and frequency to enter (**p<0.01) the open arm of the elevated plus maze indicating an increase in anxiety-related behaviors (Figure 5B and C, Student’s t-test). However, these animals did not exhibit anxiety behaviors in the open field assay or overall locomotor activity (Figure 5D - F). GPR83 KD in the CeA or NAc had no effect on these behaviors except for a decrease in the frequency to enter the open arm of the elevated plus maze in the case of the NAc (Figure 5G-R). Overall, these data indicate that GPR83 expression in the BLA regulates anxiety levels in female mice however, revealing these differences depends on the sensitivity of the assay used.

**Figure 5:**
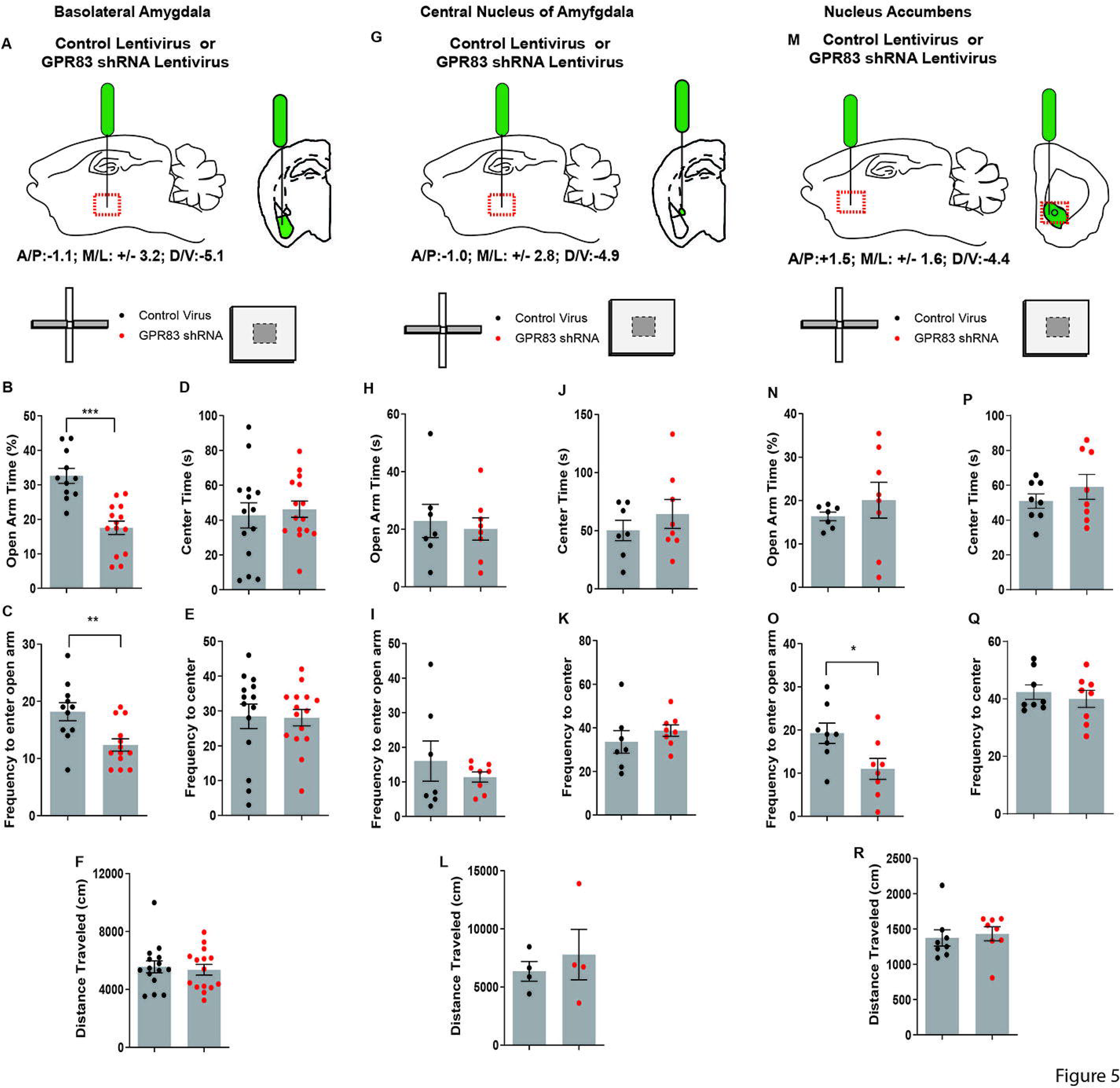
GPR83 knockdown in the BLA, but not the CeA or NAc increases anxiety-related behaviors in female mice. Schematic of injection of control or GPR83 shRNA lentivirus into the **(A)** BLA, **(G)** CeA or **(M)** nucleus accumbens. Effect of brain region specific GPR83 knockdown in mice on open arm time in the elevated plus maze **(B, H, N)** and on the frequency to enter the open arm **(C, I, O)**. Effect of brain region specific GPR83 knockdown in mice on the center time in the open field assay **(D, J, P)**, on frequency to enter the center **(E, K, Q)** and on the distance traveled **(F, L, R)**. Data are represented as mean ± SEM and analyzed using Student’s T-test, % =open arm time/ (open arm + closed arm time), *p<0.05, **p<0.01, ***p<0.001, BLA, CV n= 11, GPR83 shRNA n=14; CeA, CV n= 7, GPR83 shRNA n=8; NAc, CV n=8, GPR84 shRNA n=8.

Next we examined the estrus cycle-dependency on anxiety following GPR83 KD in the BLA. For this, we monitored the estrus cycle by taking vaginal swabs immediately following behavioral testing which were categorized by two blind observers into the different stages by the presence of leukocytes, nucleated and cornified epithelial cells. Oestrus was defined as mice in the proestrus and estrus phase during which circulating hormone levels peak. Diestrus was defined as mice in metestrus and diestrus during which circulating hormone levels are lower (Wood et al., 2007; Miller and Takahashi, 2014). This analysis revealed that both oestrus and diestrus mice with GPR83 KD in the BLA exhibit significant decreases in time spent in the open arms indicating that the increases in anxiety are not estrus cycle-dependent (Figure 6B; 2-way ANOVA; Interaction F_(1,21)_=0.75, p=0.3955; GPR83 KD F_(1,21)_=22.83, p=0.0.0001; Estrus cycle F_(1,21)_=0.01, p=0.9314; Bonferroni post-hoc test, Oestrus Control virus vs GPR83 KD, p<0.0001; Diestrus Control virus vs GPR83 KD p<0.05). There was a trend for amygdala GPR83 KD mice in diestrus to enter the open arm less frequently than mice in oestrus (Figure 6 C; 2-way ANOVA; Interaction F_(1,23)_=0.38, p=0.5460; GPR83 KD F_(1,23)_=2.07, p=0.1649; Estrus cycle F_(1,23)_=1.50, p=0.2328; Student’s test, Diestrus Control virus vs GPR83 shRNA, p<0.1272). Finally, further analysis of estrus cycle effects reveals a possible effect of estrus cycle on center time and frequency to enter the center which may be explained by an overall decrease in activity of mice in diestrus revealed by decreases in locomotor activity (Figure 6D-F; 2-way ANOVA; Interaction F_(1,26)_=0.02, p=0.9023; GPR83 KD F_(1,26)_=0.00, p=0.9668; Estrus cycle F_(1,26)_=3.94, p=0.0577; Student’s test Oestrus vs Diestrus, # p<0.05). Together, these data indicate that there is no difference in anxiety behaviors between mice in oestrus vs diestrus however, diestrus decreases the overall activity levels of mice.

**Figure 6:**
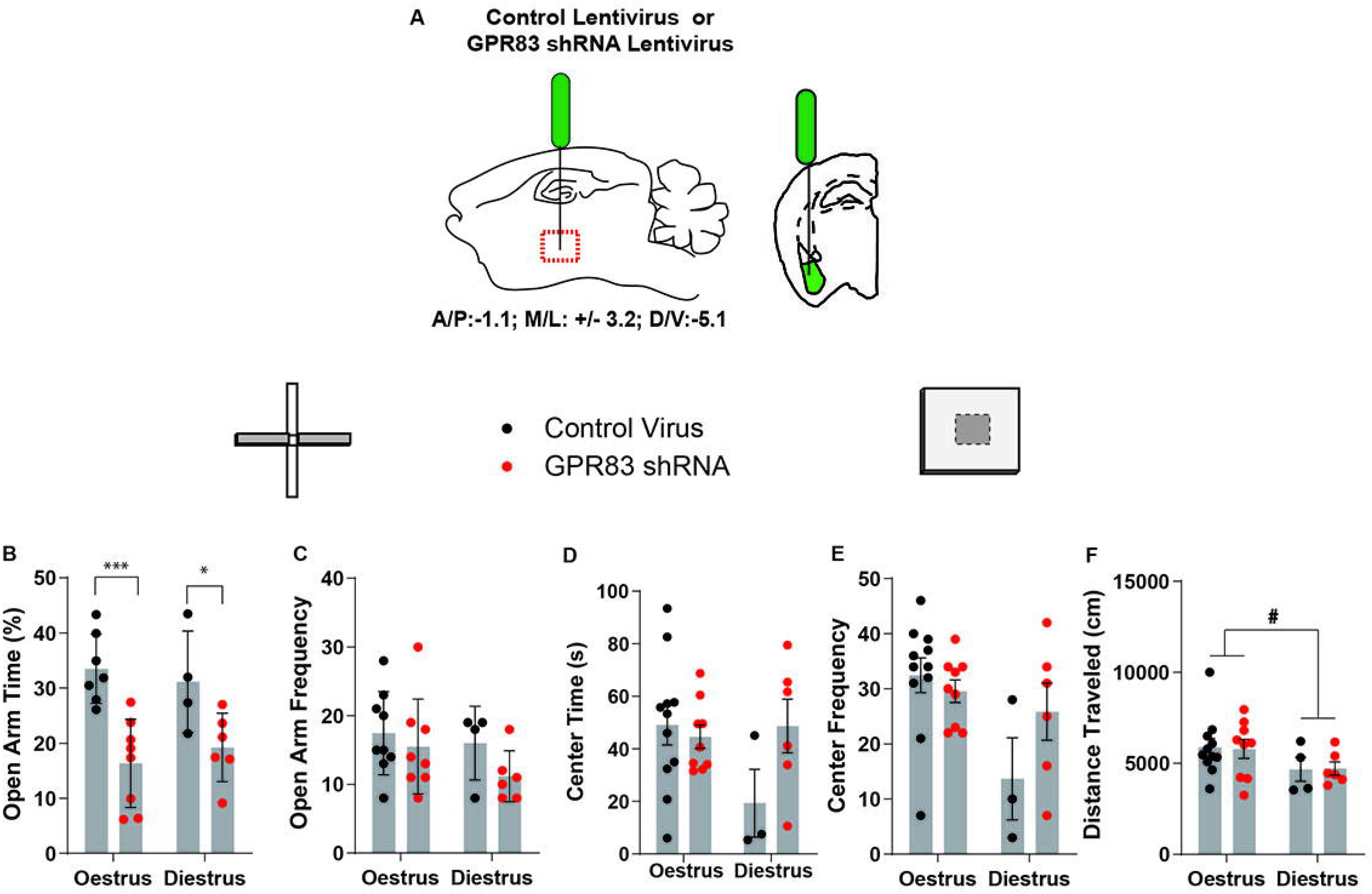
Analysis of estrus cycle-dependent differences in anxiety-related behaviors following GPR83 knockdown in the BLA. **(A)** Schematic of injection of control or GPR83 shRNA lentivirus into the BLA. Analysis of estrus cycle-dependent differences following GPR83 knockdown in the BLA, on the elevated plus maze and open field assays, measuring open arm time **(B)**, frequency to enter the open arm **(C),** center time **(D)**, frequency to enter the center **(E)** and distance traveled **(F)**. Data are represented as mean ± SEM and analyzed using 2-way ANOVA following Bonferroni’s post-hoc test, %= open arm time/ (open arm + closed arm time),*p<0.05, ***p<0.001, Student’s t-test, #p<0.05; n=3-11 per group.

## Discussion

Early studies have shown that GPR83 expression is regulated by the glucocorticoid receptor agonist dexamethasone (Harrigan et al., 1989; Adams et al., 2003), suggesting a role for GPR83 in stress and anxiety responses, since glucocorticoid release is a hallmark of the stress response. In fact, studies have reported that mice lacking GPR83 are resistant to stress-induced anxiety (Vollmer et al., 2013). However, these studies did not examine sex-differences or the specific brain regions where GPR83-mediated regulation of anxiety-related behavior may occur.

We found that global loss of GPR83 leads to a decrease in anxiety-related behaviors which is more prominent in male compared to female mice. In agreement with other studies (Simpson et al., 2012), we found that female wild type mice tend to display lower baseline levels of anxiety; this could account for the lack of effect of the global GPR83 KO on anxiety-related behaviors in female mice. In fact, these studies found that female mice were resistant to treatment with the anti-anxiety drug, diazepam, as compared to males, likely due to a floor effect in the female mice (Simpson et al., 2012). These data support the concept that lower baseline levels of anxiety in female mice may be a significant factor in screening treatments for anxiety. Together these data highlight the importance of examining the effectiveness of anxiety treatments on both males and females in preclinical models using multiple behavioral assays of anxiety, since specific assays may not be ideal for both sexes. In this context, future in-depth characterization of the role of GPR83 in anxiety will require screening in alternate assays besides the ones described in this study (elevated plus maze, open field), such as novelty suppressed feeding, marble burying etc. in order to fully understand the role of this receptor system in modulating nuances of anxiety behaviors.

There are several classes of drugs for the treatment of anxiety disorders including those that act to control the balance between GABA and glutamate transmission (Murrough et al., 2015). The BLA contains local inhibitory neurons that regulate excitatory projections to the CeA, ventral hippocampus (vHPC), medial prefrontal cortex (mPFC), bed nucleus of the stria terminalis (BNST), and NAc (Sah et al., 2003; Janak and Tye, 2015b; Tovote et al., 2015). While activation of the BLA projection to the mPFC and vHPC induces anxiety-related behaviors, activation of BLA projection to the CeA and BNST results in anxiolysis (Nascimento Häckl and Carobrez, 2007; Tye et al., 2011; Felix-Ortiz et al., 2013, 2016; Kim et al., 2013; Felix-Ortiz and Tye, 2014; Lowery-Gionta et al., 2018). Moreover, the circuits from the BLA to NAc and the BLA to CeA have been shown to encode positive and negative valence, respectively (Stuber et al., 2011; Namburi et al., 2015; Beyeler et al., 2016), suggesting that a complex network of circuits contribute to the overall anxiety state.

In our studies, complete removal of GPR83 in the knockout animal produced decreases in anxiety- related behaviors while specific knockdown in the BLA resulted in more anxiety-related behaviors. One reason for this discrepancy between the effect of global knockout versus local knockdown in the BLA may be due to an imbalance in these outgoing circuitries, suggesting the GPR83 tone from BLA contributes more to the anxiolysis, since there is an increase in anxiety with loss of GPR83 in this region, while global KO of GPR83 expression offsets this change in amygdalar tone. Previous studies have detected GPR83 expression in the PFC, hypothalamus, NAc, hippocampus and BNST (Pesini et al., 1998; Brezillon et al., 2001; Wang et al., 2001; Eberwine and Bartfai, 2011; Dubins et al., 2012; Müller et al., 2013; Lueptow et al., 2018; Fakira et al., 2019), though the role of GPR83 in each of these brain regions has yet to be explored. The current studies suggest that removing GPR83 from all these regions may shift the overall output in a direction which favors less anxiety and the mechanisms that underlie this remains to be examined.

Though reducing expression of GPR83 in the BLA uncovered a shift towards increasing anxiety-related behaviors, GPR83 KD in the CeA and NAc had little to no effect. Previous studies showed that blocking excitatory output from the BLA to the CeA or mPFC resulted in a shift towards increasing anxiety-related behaviors (Tye et al., 2011; Felix-Ortiz et al., 2016; Lowery-Gionta et al., 2018). Therefore, it is possible that GPR83 expression on interneurons in the BLA regulates inhibitory control. In line with this concept, our studies identified GPR83 expression on parvalbumin positive GABAergic neurons, which are known to form perisomatic synapses, i.e. along the soma, axon initial segment and proximal dendrites, of excitatory pyramidal neurons in the BLA representing half of their inhibitory input (McDonald and Mascagni, 2001; Muller et al., 2006). Therefore, these parvalbumin expressing neurons are in a prime position to gate output from the BLA. Furthermore, recent studies demonstrated that suppressing parvalbumin neuron activity in the BLA upregulates anxiety-related behaviors (Luo et al., 2020) similar to the increases in anxiety seen following GPR83 knockdown in the BLA. This suggests that reducing GPR83 expression on parvalbumin neurons may suppress parvalbumin neuron activity thereby resulting in a net increase in excitatory output to downstream brain regions. In order to determine the role of GPR83 on excitatory circuits in the BLA, future studies investigating the impact of GPR83 knockdown on inhibitory and excitatory neurotransmission are necessary.

Because GPR83 has been implicated in anxiety-related behaviors, and subcellular regulation of the receptor is altered by stress-related glucocorticoids, we investigated the impact of the glucocorticoid agonist, dexamethasone, on regulation of GPR83 expression in a sex and brain region specific manner. In the amygdala, we find that dexamethasone treatment reduced GPR83 expression in female mice but had no effect in males. Consistent with this a previous study using male mice reported that GPR83 expression in the amygdala was not effected by dexamethasone treatment (Adams et al., 2003). Our behavioral studies indicate divergent sex effects of GPR83 on behavior. In the NAc, we find that dexamethasone induced opposing effects on GPR83 expression, increasing expression in males while decreasing expression in females. Our observations with male mice are in contrast to those of Adams et al (2003) that reported decrease in GPR83 expression in NAc following dexamethasone treatment. This discrepancy could be due to a number of factors including the strain of mice used (C57Bl6 vs ICR), sensitivity of the technique used (in situ hybridization vs real-time qPCR), and/or the effects of oestrus cycle hormones in the female mice.

We have also found variability in GPR83 expression in individual male mice. It is known that the levels of corticosterone, the naturally occurring glucocorticoid in rodents, are higher in females compared to males (Nguyen et al., 2020). Based on this, female mice may have more stable expression of glucocorticoid regulated proteins such as GPR83 which may explain why female mice have less variable GPR83 expression. Additionally, this may explain why the effect of dexamethasone on GPR83 was more consistent between brain regions in females, where we observed a dexamethasone-induced decrease in both the NAc and amygdala. The higher levels of corticosterone in females may also contribute to the sex-differences in baseline anxiety and GPR83-mediated regulation of anxiety levels reported above.

Another important finding of this study is that knockdown of GPR83 in the BLA of female mice increased anxiety-related behaviors in the EPM test irrespective of whether the animals were in the oestrus or diestrus stage. GPR83 expression in the uterus is regulated during the estrus cycle in an estrogen and progesterone dependent manner (Parobchak et al., 2020) suggesting that circulating hormone levels may influence GPR83 function. Studies have shown that females in proestrus display decreased anxiety-related behaviors which corresponded with higher levels of progesterone and its metabolite 5α-pregnan-3α-ol-20-one (3α-5α-THP; allopregnalone) (Frye et al., 2000). In line with this, treatment of ovariectomized rats with progesterone is anxiolytic and corresponded with the potentiation of GABA_A_ receptor currents (Gulinello and Smith, 2003). Subsequent studies found that administration of allopregnalone is anxiolytic when administered acutely. However, chronic allopregnalone treatment is anxiogenic in both male and female mice and alters the anxiolytic potential of the benzodiazepine ligands lorazepam and flumazenil (Gulinello and Smith, 2003).

Our studies did not identify any differences in anxiety between wild type females in oestrus versus diestrus. This may be because in our studies we had pooled mice in groups with high circulating hormones (oestrus- proestrus/estrus) and low circulating hormones (diestrus-metestrus/diestrus) (Miller and Takahashi, 2014). By pooling together animals with varying levels of individual hormones, these differences in anxiety levels may have reached below detectable threshold. Given the role of progesterone/allopregnalone in modulating anxiety-related behaviors via regulation of GABAergic function, and that GPR83 expression is regulated by estrogen and progesterone in the uterus (Parobchak et al., 2020) further studies are needed to determine if there is a relationship between GPR83 and progesterone levels in the brain.

In summary, our studies suggest that GPR83 is differentially regulated between male and female mice. Furthermore, regional changes in expression of GPR83 significantly impacts the overall tone of anxiety-related circuitry, and specifically, GPR83 expression in the BLA may be a primary output node for regulating anxiety-related behavior.

## Acknowledgements

The authors wish to thank Andrei Jeltyi for assistance in genotyping and animal husbandry and Dr. Ivone Gomes for critical reading of the manuscript. This work was supported by NIH grants R01-DA008863 and R01-NS026880 (to LAD). Authors have no conflict of interest.

## References

Adams, F., Grassie, M., Shahid, M., Hill, D. R., and Henry, B. (2003). Acute oral dexamethasone administration reduces levels of orphan GPCR glucocorticoid-induced receptor (GIR) mRNA in rodent brain: Potential role in HPA-axis function. Mol. Brain Res. 117, 39–46. doi:10.1016/S0169-328X(03)00280-8.

Babaev, O., Piletti Chatain, C., and Krueger-Burg, D. (2018). Inhibition in the amygdala anxiety circuitry. Exp. Mol. Med. doi:10.1038/s12276-018-0063-8.

Berezniuk, I., Rodriguiz, R. M., Zee, M. L., Marcus, D. J., Pintar, J., Morgan, D. J., et al. (2017). ProSAAS-derived peptides are regulated by cocaine and are required for sensitization to the locomotor effects of cocaine. J. Neurochem. doi:10.1111/jnc.14209.

Beyeler, A., Namburi, P., Glober, G. F., Simonnet, C., Calhoon, G. G., Conyers, G. F., et al. (2016). Divergent Routing of Positive and Negative Information from the Amygdala during Memory Retrieval. Neuron. doi:10.1016/j.neuron.2016.03.004.

Bobeck, E. N., Gomes, I., Pena, D., Cummings, K. A., Clem, R. L., Mezei, M., et al. (2017). The BigLEN-GPR171 peptide receptor system within the basolateral amygdala regulates anxiety-like behavior and contextual fear conditioning. Neuropsychopharmacology. doi:10.1038/npp.2017.79.

Boivin, J. R., Piekarski, D. J., Wahlberg, J. K., and Wilbrecht, L. (2017). Age, sex, and gonadal hormones differently influence anxiety- and depression-related behavior during puberty in mice. Psychoneuroendocrinology. doi:10.1016/j.psyneuen.2017.08.009.

Brezillon, S., Detheux, M., Parmentier, M., Hokfelt, T., and Hurd, Y. L. (2001). Distribution of an orphan G-protein coupled receptor (JP05) mRNA in the human brain. Brain Res. 921, 21–30. doi:S0006-8993(01)03068-2 [pii].

Dubins, J. S., Sanchez-Alavez, M., Zhukov, V., Sanchez-Gonzalez, A., Moroncini, G., Carvajal-Gonzalez, S., et al. (2012). Downregulation of GPR83 in the hypothalamic preoptic area reduces core body temperature and elevates circulating levels of adiponectin. Metabolism. 61, 1486–1493. doi:10.1016/j.metabol.2012.03.015.

Eberwine, J., and Bartfai, T. (2011). Single cell transcriptomics of hypothalamic warm sensitive neurons that control core body temperature and fever response: Signaling asymmetry and an extension of chemical neuroanatomy. Pharmacol. Ther. 129, 241–259. doi:10.1016/j.pharmthera.2010.09.010.

Enterría-Morales, D., del Rey, N. L.-G., Blesa, J., López-López, I., Gallet, S., Prévot, V., et al. (2020). Molecular targets for endogenous glial cell line-derived neurotrophic factor modulation in striatal parvalbumin interneurons. Brain Commun. doi:10.1093/braincomms/fcaa105.

Fakira, A. K., Peck, E. G., Liu, Y., Lueptow, L. M., Trimbake, N. A., Han, M. H., et al. (2019). The role of the neuropeptide PEN receptor, GPR83, in the reward pathway: Relationship to sex-differences. Neuropharmacology. doi:10.1016/j.neuropharm.2019.107666.

Felix-Ortiz, A. C., Beyeler, A., Seo, C., Leppla, C. A., Wildes, C. P., and Tye, K. M. (2013). BLA to vHPC inputs modulate anxiety-related behaviors. Neuron. doi:10.1016/j.neuron.2013.06.016.

Felix-Ortiz, A. C., Burgos-Robles, A., Bhagat, N. D., Leppla, C. A., and Tye, K. M. (2016). Bidirectional modulation of anxiety-related and social behaviors by amygdala projections to the medial prefrontal cortex. Neuroscience. doi:10.1016/j.neuroscience.2015.07.041.

Felix-Ortiz, A. C., and Tye, K. M. (2014). Amygdala inputs to the ventral hippocampus bidirectionally modulate social behavior. J. Neurosci. doi:10.1523/JNEUROSCI.4257-13.2014.

Foster, S. R., Hauser, A. S., Vedel, L., Strachan, R. T., Huang, X. P., Gavin, A. C., et al. (2019). Discovery of Human Signaling Systems: Pairing Peptides to G Protein-Coupled Receptors. Cell. doi:10.1016/j.cell.2019.10.010.

Fricker, L. D., McKinzie, A. A., Sun, J., Curran, E., Qian, Y., Yan, L., et al. (2000). Identification and characterization of proSAAS, a granin-like neuroendocrine peptide precursor that inhibits prohormone processing. J. Neurosci. doi:10.1523/jneurosci.20-02-00639.2000.

Frye, C. A., Petralia, S. M., and Rhodes, M. E. (2000). Estrous cycle and sex differences in performance on anxiety tasks coincide with increases in hippocampal progesterone and 3α,5α-THP. Pharmacol. Biochem. Behav. doi:10.1016/S0091-3057(00)00392-0.

Gilpin, N. W., Herman, M. A., and Roberto, M. (2015). The Central Amygdala as an Integrative Hub for Anxiety and Alcohol Use Disorders. Biol. Psychiatry 77, 859–869. doi:10.1016/j.biopsych.2014.09.008.

Gomes, I., Bobeck, E. N., Margolis, E. B., Gupta, A., Sierra, S., Fakira, A. K., et al. (2016). Identification of GPR83 as the receptor for the neuroendocrine peptide PEN. Sci. Signal. 9. doi:10.1126/scisignal.aad0694.

Griebel, G., and Holmes, A. (2013). 50 years of hurdles and hope in anxiolytic drug discovery. Nat. Rev. Drug Discov. doi:10.1038/nrd4075.

Gulinello, M., and Smith, S. S. (2003). Anxiogenic effects of neurosteroid exposure: Sex differences and altered GABAA receptor pharmacology in adult rats. J. Pharmacol. Exp. Ther. doi:10.1124/jpet.102.045120.

Harrigan, M. T., Baughman, G., Campbell, N. F., and Bourgeois, S. (1989). Isolation and characterization of glucocorticoids- and cyclic AMP-induced genes in T lymphocytes. Mol. Cell. Biol. 9, 3438–3446.

Harrigan, M. T., Campbell, N. F., and Bourgeois, S. (1991). Identification of a gene induced by glucocorticoids in murine T-cells: a potential G protein-coupled receptor. Mol.Endocrinol. 5, 1331–1338. doi:10.1210/mend-5-9-1331.

Hoshino, A., Helwig, M., Rezaei, S., Berridge, C., Eriksen, J. L., and Lindberg, I. (2014). A novel function for proSAAS as an amyloid anti-aggregant in Alzheimer’s disease. J. Neurochem. 128, 419–430. doi:10.1111/jnc.12454.

Janak, P. H., and Tye, K. M. (2015a). From circuits to behaviour in the amygdala. Nature 517, 284–292. doi:10.1038/nature14188.

Janak, P. H., and Tye, K. M. (2015b). From circuits to behaviour in the amygdala. Nature. doi:10.1038/nature14188.

Kim, S. Y., Adhikari, A., Lee, S. Y., Marshel, J. H., Kim, C. K., Mallory, C. S., et al. (2013). Diverging neural pathways assemble a behavioural state from separable features in anxiety. Nature. doi:10.1038/nature12018.

Lowery-Gionta, E. G., Crowley, N. A., Bukalo, O., Silverstein, S., Holmes, A., and Kash, T. L. (2018). Chronic stress dysregulates amygdalar output to the prefrontal cortex. Neuropharmacology. doi:10.1016/j.neuropharm.2018.06.032.

Lu, L. F., Gavin, M. A., Rasmussen, J. P., and Rudensky, A. Y. (2007). G protein-coupled receptor 83 is dispensable for the development and function of regulatory T cells. Mol Cell Biol 27, 8065–8072. doi:10.1128/MCB.01075-07.

Lueptow, L. M., Devi, L. A., and Fakira, A. K. (2018). Targeting the Recently Deorphanized Receptor GPR83 for the Treatment of Immunological, Neuroendocrine and Neuropsychiatric Disorders. Prog. Mol. Biol. Transl. Sci. doi:10.1016/bs.pmbts.2018.07.002.

Luo, Z. Y., Huang, L., Lin, S., Yin, Y. N., Jie, W., Hu, N. Y., et al. (2020). Erbin in Amygdala Parvalbumin-Positive Neurons Modulates Anxiety-like Behaviors. Biol. Psychiatry. doi:10.1016/j.biopsych.2019.10.021.

Mack, S. M., Gomes, I., and Devi, L. A. (2019). Neuropeptide PEN and Its Receptor GPR83: Distribution, Signaling, and Regulation. ACS Chem. Neurosci. doi:10.1021/acschemneuro.8b00559.

McDonald, A. J., and Mascagni, F. (2001). Colocalization of calcium-binding proteins and GABA in neurons of the rat basolateral amygdala. Neuroscience. doi:10.1016/S0306-4522(01)00214-7.

McLean, A. C., Valenzuela, N., Fai, S., and Bennett, S. A. L. (2012). Performing vaginal lavage, crystal violet staining, and vaginal cytological evaluation for mouse estrous cycle staging identification. J. Vis. Exp. doi:10.3791/4389.

Miller, B. H., and Takahashi, J. S. (2014). Central circadian control of female reproductive function. Front. Endocrinol. (Lausanne). doi:10.3389/fendo.2013.00195.

Morgan, D. J., Wei, S., Gomes, I., Czyzyk, T., Mzhavia, N., Pan, H., et al. (2010). The propeptide precursor proSAAS is involved in fetal neuropeptide processing and body weight regulation. J. Neurochem. 113, 1275–1284. doi:10.1111/j.1471-4159.2010.06706.x.

Muller, J. F., Mascagni, F., and McDonald, A. J. (2006). Pyramidal cells of the rat basolateral amygdala: Synaptology and innervation by parvalbumin-immunoreactive interneurons. J. Comp. Neurol. doi:10.1002/cne.20832.

Müller, T. D., Müller, A., Yi, C. X., Habegger, K. M., Meyer, C. W., Gaylinn, B. D., et al. (2013). The orphan receptor Gpr83 regulates systemic energy metabolism via ghrelin-dependent and ghrelin-independent mechanisms. Nat. Commun. 4. doi:10.1038/ncomms2968.

Murrough, J. W., Yaqubi, S., Sayed, S., and Charney, D. S. (2015). Emerging drugs for the treatment of anxiety. Expert Opin. Emerg. Drugs. doi:10.1517/14728214.2015.1049996.

Mzhavia, N., Qian, Y., Feng, Y., Che, F., Devi, L., and Fricker, L. (2002). Processing of proSAAS in neuroendocrine cell lines. Biochem. J 76, 67–76. doi:10.1042/0264-6021:3610067.

Namburi, P., Beyeler, A., Yorozu, S., Calhoon, G. G., Halbert, S. A., Wichmann, R., et al. (2015). A circuit mechanism for differentiating positive and negative associations. Nature. doi:10.1038/nature14366.

Nascimento Häckl, L. P., and Carobrez, A. P. (2007). Distinct ventral and dorsal hippocampus AP5 anxiolytic effects revealed in the elevated plus-maze task in rats. Neurobiol. Learn. Mem. doi:10.1016/j.nlm.2007.04.007.

Nguyen, K., Kanamori, K., Shin, C. S., Hamid, A., and Lutfy, K. (2020). The impact of sex on changes in plasma corticosterone and cotinine levels induced by nicotine in c57bl/6j mice. Brain Sci. doi:10.3390/brainsci10100705.

Palanza, P. (2001). Animal models of anxiety and depression: How are females different? Neurosci. Biobehav. Rev. doi:10.1016/S0149-7634(01)00010-0.

Parobchak, N., Rao, S., Negron, A., Schaefer, J., Bhattacharya, M., Radovick, S., et al. (2020). Uterine Gpr83 mRNA is highly expressed during early pregnancy and GPR83 mediates the actions of PEN in endometrial and non-endometrial cells. F&S Sci. doi:10.1016/j.xfss.2020.06.001.

Paxinos, G., and Franklin, K. B. J. (2012). Paxinos and Franklin’s the Mouse Brain in Stereotaxic Coordinates.

Pesini, P., Detheux, M., Parmentier, M., and Hökfelt, T. (1998). Distribution of a glucocorticoid-induced orphan receptor (JP05) mRNA in the central nervous system of the mouse. Mol. Brain Res. 57, 281–300. doi:10.1016/S0169-328X(98)00099-0.

Ravenelle, R., Neugebauer, N. M., Niedzielak, T., and Donaldson, S. T. (2014). Sex differences in diazepam effects and parvalbumin-positive GABA neurons in trait anxiety Long Evans rats. Behav. Brain Res. doi:10.1016/j.bbr.2014.04.048.

Sah, P., Faber, E. S. L., De Armentia, M. L., and Power, J. (2003). The amygdaloid complex: Anatomy and physiology. Physiol. Rev. doi:10.1152/physrev.00002.2003.

Simpson, J., Ryan, C., Curley, A., Mulcaire, J., and Kelly, J. P. (2012). Sex differences in baseline and drug-induced behavioural responses in classical behavioural tests. Prog. Neuro-Psychopharmacology Biol. Psychiatry. doi:10.1016/j.pnpbp.2012.02.004.

Stuber, G. D., Sparta, D. R., Stamatakis, A. M., Van Leeuwen, W. A., Hardjoprajitno, J. E., Cho, S., et al. (2011). Excitatory transmission from the amygdala to nucleus accumbens facilitates reward seeking. Nature 475, 377–382. doi:10.1038/nature10194.

Tovote, P., Fadok, J. P., and Lüthi, A. (2015). Neuronal circuits for fear and anxiety. Nat. Rev. Neurosci. doi:10.1038/nrn3945.

Tye, K. M., Prakash, R., Kim, S. Y., Fenno, L. E., Grosenick, L., Zarabi, H., et al. (2011). Amygdala circuitry mediating reversible and bidirectional control of anxiety. Nature. doi:10.1038/nature09820.

Urakawa, S., Takamoto, K., Hori, E., Sakai, N., Ono, T., and Nishijo, H. (2013). Rearing in enriched environment increases parvalbumin-positive small neurons in the amygdala and decreases anxiety-like behavior of male rats. BMC Neurosci. doi:10.1186/1471-2202-14-13.

Vollmer, L., Ghosal, S., A Rush, J., R Sallee, F., P Herman, J., Weinert, M., et al. (2013). Attenuated stress-evoked anxiety, increased sucrose preference and delayed spatial learning in glucocorticoid-induced receptor-deficient mice. Genes. Brain. Behav. 12, 241–9. doi:10.1111/j.1601-183X.2012.00867.x.

Wang, J., Cunningham, R., Zetterberg, H., Asthana, S., Carlsson, C., Okonkwo, O., et al. (2016a). Label-free quantitative comparison of cerebrospinal fluid glycoproteins and endogenous peptides in subjects with Alzheimer’s disease, mild cognitive impairment, and healthy individuals. Proteomics - Clin. Appl. doi:10.1002/prca.201600009.

Wang, D., J.P., Herman, L.M., Pritchard, R.H., Spitzer, R.L., Ahlbrand, G.L., Kramer, et al. (2001). Cloning, expression and regulation of a glucocorticoid-induced receptor in rat brain: effect of repetitive amphetamine. J. Neurosci. 21, 9027–9035.

Wang, L., Shen, M., Jiang, C., Ma, L., and Wang, F. (2016b). Parvalbumin interneurons of central amygdala regulate the negative affective states and the expression of corticotrophin-releasing hormone during morphine withdrawal. Int. J. Neuropsychopharmacol. doi:10.1093/ijnp/pyw060.

Wardman, J. H., Berezniuk, I., Di, S., Tasker, J. G., and Fricker, L. D. (2011). ProSAAS-derived peptides are colocalized with neuropeptide Y and function as neuropeptides in the regulation of food intake. PLoS One 6. doi:10.1371/journal.pone.0028152.

Wei, S., Feng, Y., Che, F.-Y., Pan, H., Mzhavia, N., Devi, L. A., et al. (2004). Obesity and diabetes in transgenic mice expressing proSAAS. J. Endocrinol. 180. doi:10.1677/joe.0.1800357.

Wood, G. A., Fata, J. E., Watson, K. L. M., and Khokha, R. (2007). Circulating hormones and estrous stage predict cellular and stromal remodeling in murine uterus. Reproduction. doi:10.1530/REP-06-0302.

Xiao, Q., Zhou, X., Wei, P., Xie, L., Han, Y., Wang, J., et al. (2020). A new GABAergic somatostatin projection from the BNST onto accumbal parvalbumin neurons controls anxiety. Mol. Psychiatry. doi:10.1038/s41380-020-0816-3.

Zhu, Z., Wang, G., Ma, K., Cui, S., and Wang, J.-H. (2017). GABAergic neurons in nucleus accumbens are correlated to resilience and vulnerability to chronic stress for major depression. Oncotarget. doi:10.18632/oncotarget.16411.

Zuloaga, D. G., Heck, A. L., De Guzman, R. M., and Handa, R. J. (2020). Roles for androgens in mediating the sex differences of neuroendocrine and behavioral stress responses. Biol. Sex Differ. doi:10.1186/s13293-020-00319-2.

